# Present and Future Ecological Niche Modeling of Rift Valley fever in East Africa in Response to Climate Change

**DOI:** 10.1101/2021.03.03.433832

**Authors:** Caroline Muema, Boniface K. Ngarega, Elishiba Muturi, Hongping Wei, Hang Yang

**Author notes:** Authors for correspondence: **Hongping Wei; E-mail**. **Hang Yang; E-mail**.

## Abstract

Rift Valley fever (RVF) has been linked with recurrent outbreaks among humans and livestock in several parts of the globe. Predicting RVF’s habitat suitability under different climate scenarios offers vital information for developing informed management schemes. The present study evaluated the probable impacts of climate change on the distribution of RVF disease in East Africa (E. A.), using the maximum entropy (MaxEnt) model and the disease outbreak cases. Considering the potential of the spread of the disease in the East Africa region, we utilized two representative concentration pathways (RCP 4.5 and RCP 8.5) climate scenarios in the 2050s and 2070s (average for 2041-2060, and 2061-2080), respectively. All models had satisfactory AUC values of more than 0.809, which are considered excellent. Jackknife tests revealed that Bio4 (temperature seasonality), land use, and population density were the main factors influencing RVF distribution in the region. From the risk maps generated, we infer that, without regulations, this disease might establish itself across more extensive areas in the region, including most of Rwanda and Burundi. The ongoing trade between East African countries and changing climates could intensify RVF spread into new geographic extents with suitable habitats for the important zoonosis. The predicted suitable areas for RVF in eastern Kenya, southern Tanzania, and Somalia overlaps to a large extent where cattle keeping and pastoralism are highly practiced, thereby signifying the urgency to manage and control the disease. This work validates RVF outbreak cases’ effectiveness to map the disease’s distribution, thus contributing to enhanced ecological modeling and improved disease tracking and control efforts in East Africa.

## Introduction

Industrial civilization progress fosters human society’s growth (1) and poses many global challenges, including economic, energy, population, environmental challenges, and climate change (2). Climate change, defined as the gradual alteration of climatic properties or weather patterns in a statistically identifiable manner, may occur naturally or be mainly anthropogenic (3). Human activities have been shown to have facilitated about 1.0°C of global warming (4), and it is projected that these temperatures may rise to 1.5°C by the 2050s if the present trend continues (5). This, in essence, will lead to cycles of extreme weather, heatwaves, and flooding.

Fluctuations in climate have been related to many prevalent diseases triggered by floods and heatwaves, which alter the transmission of infectious diseases (6–7). Diseases transmitted by vectors have a propensity for association with specific environmental conditions, following their sensitivity to those conditions (8–9). The vector limit of climate tolerance constrains the occurrence and distribution of viral zoonosis (5, 10). Indeed, climate change will likely affect vector-borne disease (VBD) distribution and their emergence in areas where the diseases have not been observed before due to human movement, animal transportation, and vector range expansion (11–12). Climate and weather may also influence pathogens’ ecology as pathogens replicate at ambient temperatures (13).

Rift Valley fever (RVF), which has intricated multispecies epidemiology, is a mosquito-borne zoonosis caused by the Rift Valley fever virus (RVFV). Previous studies have revealed that many vectors are responsible for RVFV transmission in Africa, including *Culex sp., Aedes sp*., and *Mansoni sp*. (14–15). RVF disease is known to cause recurrent epidemics in Africa, especially in the East African region (16–17), where it has led to significant economic losses due to livestock deaths and human illness (18).

The extent and frequency of weather events are projected to change with global climate change (19), which will, in turn, increase the risk, frequency, and distribution of RVF (18, 20). Climate variations in areas with similar conditions could result in new RVF outbreaks and overburden in RVF endemic regions (21–22). Besides, future climate change effects have proven to be challenging to evaluate, but infectious zoonotic diseases will undoubtedly have an increased impact on global health (7). Therefore, improved mitigation and management strategies of climate-sensitive zoonosis are unreservedly essential (23), and the effective modeling of the climate change impacts on their spread is vital for understanding the disease and will help future interventions (24).

Previous studies have utilized different modeling approaches to determine the influence of climate change on RVF distribution. For instance, the SEIR model has been used to assess the climate change impact on RVF dynamics by combining both vertical transmission and human to mosquito transmission with climate-driven parameters, i.e., temperature and precipitation (21). Mathematical models, e.g., the Liverpool RVF model, have been used to describe the intricate relationship between the host epidemiology, the vector’s life cycle coupled with climatic factors (25). Ecoepidemiological mechanistic models have been utilized to explain how vectors’ ecology and RVF’s epidemiology are influenced by temperature and the presence of water bodies (15). Bayesian models have been implemented to produce risk maps of RVF using spatial and seasonal environmental drivers, together with the effects of climatic oscillations (18). Predictive models, e.g., the Species distribution models (SDMs), commonly reffered to as Ecological Niche Models (ENMs), have been applied to assess the distribution of RVF in Africa using environmental covariates; either regionally (26) or locally (27–29).

The fine-scale assessment of RVF disease outbreak and distribution in East Africa in an ENM approach has been scantly assessed under a climate change context (17), and limited information is available on the influence of global climate change on RVF distribution under future projections. It has otherwise been overscored by the overall modeling of the disease’s distribution across the African continent reported elsewhere (18). Therefore, we build upon the previous studies framework to bridge this knowledge gap by conducting the fine-scale distribution of RVF using environmental covariates (i.e., bioclimatic, land-use, livestock, and human density variables) to assess their influence on the distribution of RVF. Here, we used the Maximum Entropy (MaxEnt) in an ENM approach to evaluate the RVF current range dynamics, identified potential hotspots for the emergence and re-emergence, predicted its future climatic suitability, and assessed the potential spread risks of RVF in East Africa. The standard and efficiency of mitigation strategies against RVF in East Africa are vital in light of the risks posed by the spread of this important zoonotic.

We (a) tested the advantages of integrating RVFV outbreaks, livestock, and human population densities in ENMs to locate areas of high environmental suitability for the disease in East Africa, and (b) generated a risk map of the suitable RVF niches. The present results highlight the importance of integrating outbreak cases to simulate zoonotic habitat suitability and assess a restricted region compared to a broader one by providing the potentially missed information or risk areas that could be otherwise overlooked, thus refining the appropriateness of ENMs.

## Results

### RVF risk distribution

We retained 233 RVF outbreak records after initial data cleaning and 186 after a spatial rarefying approach (Figure 1). Package ENmeval results submitted the usage of hinge features for RVFV modeling. A model is good when with AUC values above 0.8. The current distribution model for RVF had an average AUC of 0.886, signifying the model’s good performance (Figure 2).

**Figure 1.**
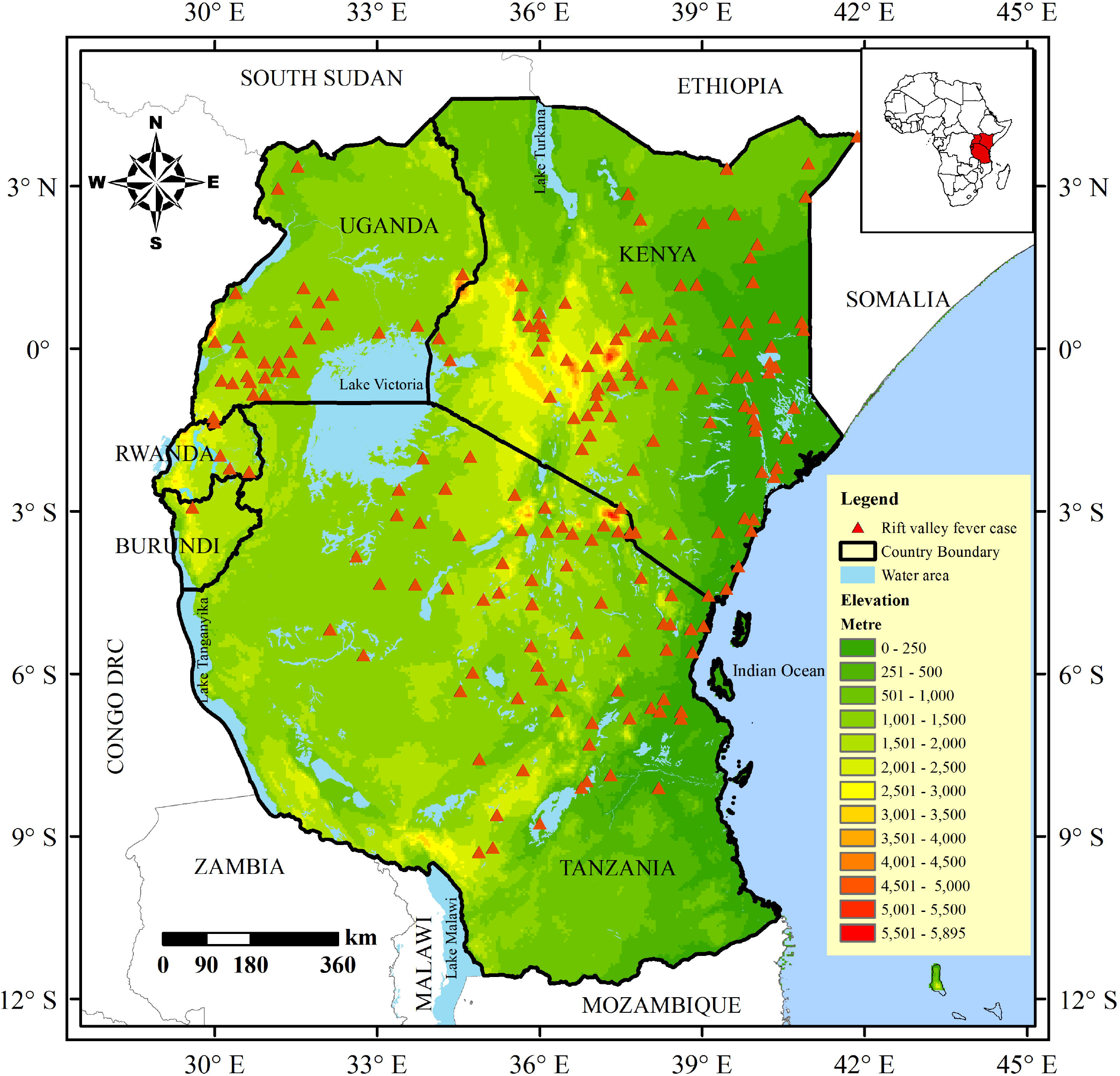
Distribution of the Rift Valley fever outbreak cases and geographic positions of occurrence points included in the modeling with elevation data.

**Figure 2.**
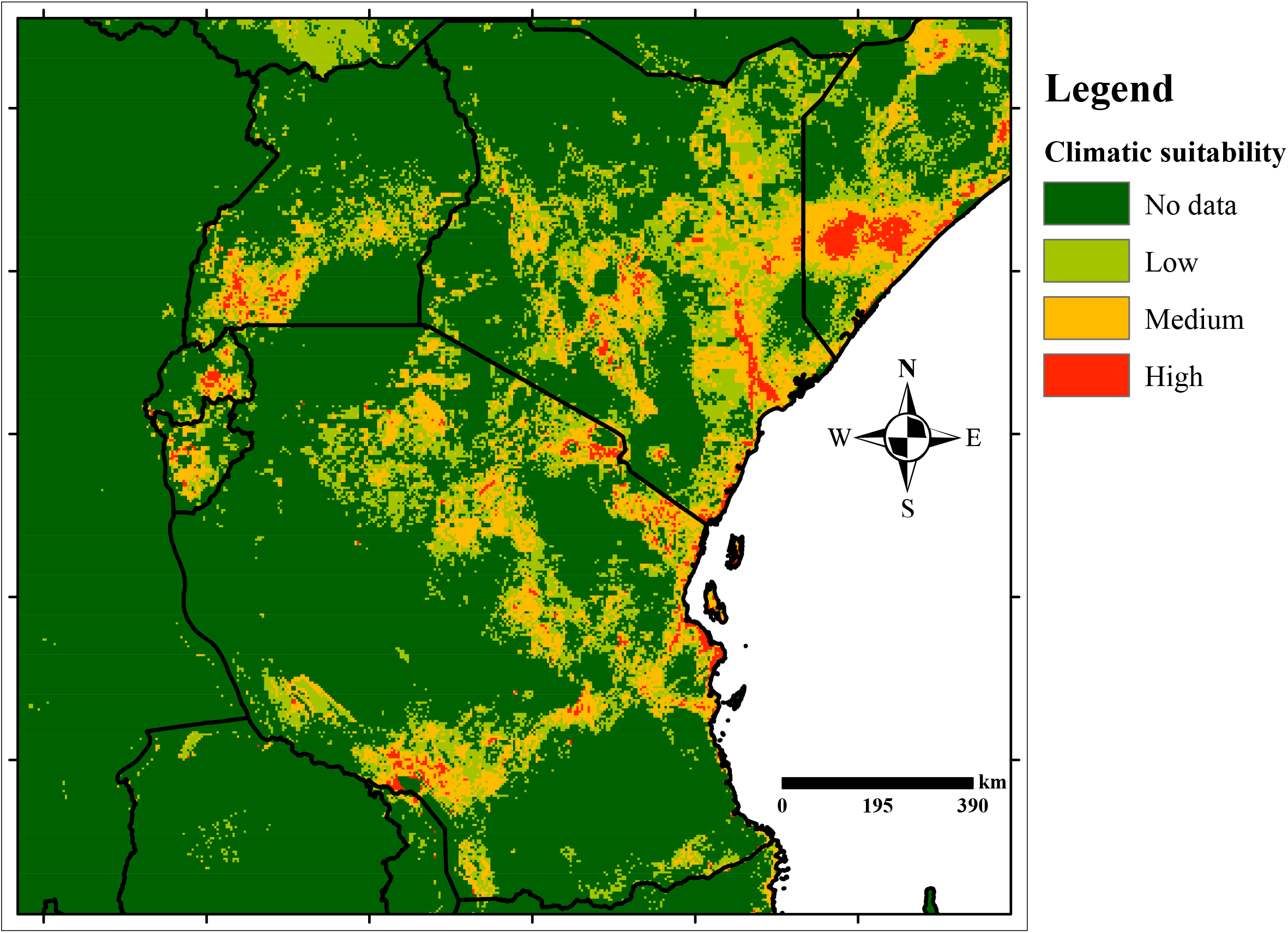
Risk map for Rift Valley fever transmission in East Africa generated after MaxEnt modeling of the current period.

Significant predictors of RVF risk distribution were temperature seasonality (Bio4), land use, population density, precipitation of coldest quarter (Bio19), elevation, and cattle, which cumulatively contributed to 55.3% of the distribution of RVF in East Africa. Of the six factors, we observe that temperature seasonality has the highest contribution (10.6%) to the final model (AUC = 0.886) (Table 2; Figure S1). Jackknife analysis showed that cattle density, sheep density, goat density, and human population density had the most significant gain when used alone, revealing that each has valuable information when used independently. When the sheep density variable is excluded, we observe that the most gain is decreased, inferring that this variable has a lot of information absent in the other variables (Figure S2). The mean temperature of the wettest quarter (Bio8) and goats were marginally important in determining the distribution of RVF in East Africa, and excluding them did not rigorously influence the performance of the model. The model predictions for the current potential distribution for the RVF were high in Kenya and Tanzania. We also identified that high RVF suitability areas concur with low elevation areas 10 - 1500 m a.s.l. (Figures 1 and 2).

**Table 1.**
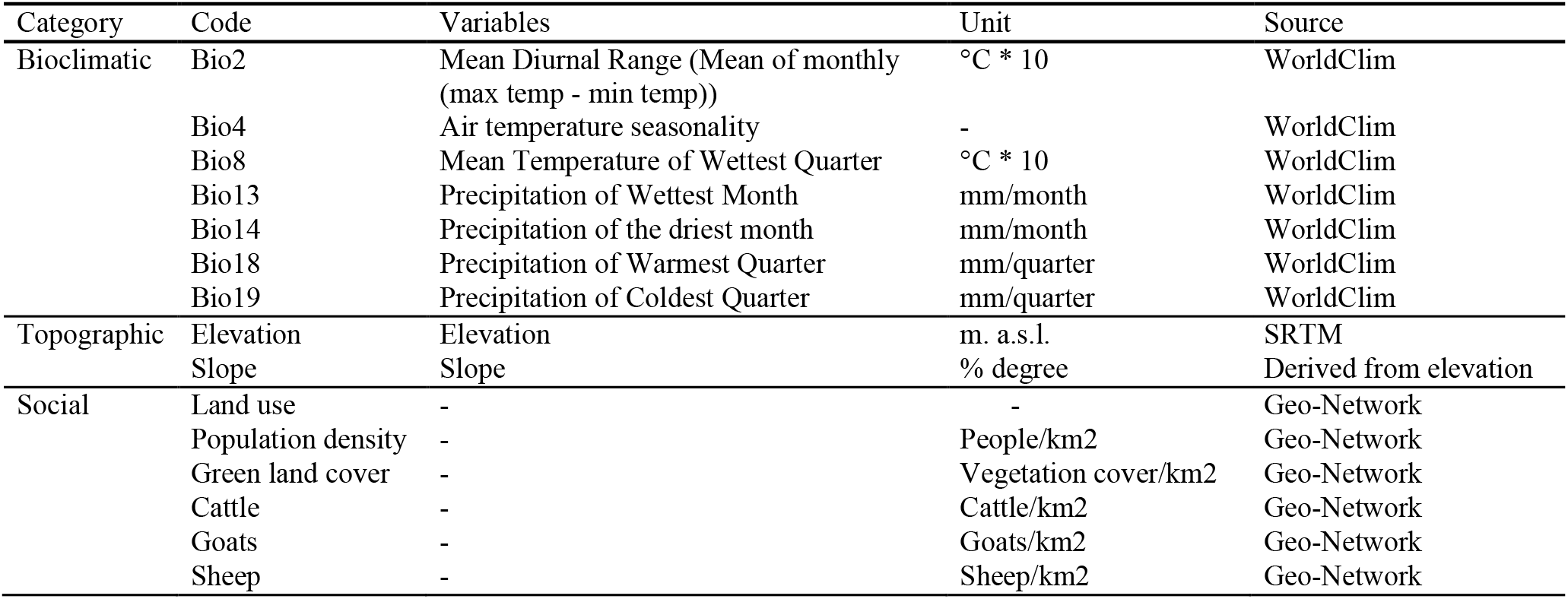
Environmental variables selected for modeling the distribution of Rift Valley fever in East Africa using MaxEnt

| Category | Code | Variables | Unit | Source |
| --- | --- | --- | --- | --- |
| Bioclimatic | Bio2 | Mean Diurnal Range (Mean of monthly (max temp - min temp)) | °C * 10 | WorldClim |
|  | Bio4 | Air temperature seasonality | - | WorldClim |
|  | Bio8 | Mean Temperature of Wettest Quarter | °C * 10 | WorldClim |
|  | Bio13 | Precipitation of Wettest Month | mm/month | WorldClim |
|  | Bio14 | Precipitation of the driest month | mm/month | WorldClim |
|  | Bio18 | Precipitation of Warmest Quarter | mm/quarter | WorldClim |
|  | Bio19 | Precipitation of Coldest Quarter | mm/quarter | WorldClim |
| Topographic | Elevation | Elevation | m. a.s.l. | SRTM |
|  | Slope | Slope | % degree | Derived from elevation |
| Social | Land use | - | - | Geo-Network |
|  | Population density | - | People/km2 | Geo-Network |
|  | Green land cover | - | Vegetation cover/km2 | Geo-Network |
|  | Cattle | - | Cattle/km2 | Geo-Network |
|  | Goats | - | Goats/km2 | Geo-Network |
|  | Sheep | - | Sheep/km2 | Geo-Network |

**Table 2.** AUC statistics for the MaxEnt’s model performance for Rift Valley Fever in East Africa

| Scenario | Current | 2050s |  | 2070s |  |
| --- | --- | --- | --- | --- | --- |
|  |  | RCP 4.5 | RCP 8.5 | RCP 4.5 | RCP 8.5 |
| **AUC mean** | 0.866 | 0.895 | 0.809 | 0.867 | 0.885 |
| **Standard Deviation** | 0.009 | 0.011 | 0.147 | 0.027 | 0.016 |

**Table 3.** Relative contributions (%) and permutation importance of environmental variables to the MaxEnt model under the current conditions of the Rift Valley fever virus in East Africa.

| Variable | Percent contribution | Permutation importance |
| --- | --- | --- |
| Bio4 | **10.6** | 8.8 |
| Land use | **9.9** | 8.4 |
| Population density | **9.8** | 9.6 |
| Bio19 | 8.8 | 10.5 |
| Elevation | 8.7 | 10.1 |
| Cattle | 7.5 | 2.9 |
| Green land cover | 6.4 | 7.5 |
| Soil type | 6.3 | 4.4 |
| Bio2 | 5.0 | 3.7 |
| Slope | 5.0 | 2.8 |
| Bio14 | 4.7 | 6.5 |
| Bio13 | 4.5 | 6.8 |
| Sheep | 4.4 | 7.4 |
| Bio18 | 4.2 | 4.2 |
| Goats | 2.4 | 3.1 |
| Bio8 | 1.9 | 3.2 |

The geographic distribution of predicted habitat suitability for RVF across East Africa is defined in more detail below and outlined in Figure 2. The highest probability occurrence is estimated for eastern Kenya and southern Somalia, east-central to southern Tanzania, north of Lake Malawi, central Rwanda, southwestern Uganda, south of Mt. Kilimanjaro, and along the coastline from Somalia to Tanzania through Kenya. The lowest habitat suitability for RVF in the current period are projected for a large area of Tanzania Plains, northern Uganda, and northwestern Kenya towards Lake Turkana.

### Future predicted RVF probable distribution for 2050 and 2070

The model performance of future RVF potential distribution was assessed according to the AUC values. All the AUC values for the period 2050 and 2070 under the two RCPs gave satisfactory values, ranging between 0.809 and 0.895, the highest (Table 2). The general distribution patterns throughout East Africa between the present-day and future models outlined realistic resemblances apart from some areas. Additionally, our model projections for 2050 and 2070 revealed that RVF differs in potential risk areas among both the RCPs, with an increase and decrease in its habitat suitability (Figure 3). Under RCP 8.5 and RCP 4.5 scenarios in the period 2050 and 2070, RVF’s habitat suitability was observed to expand in the Somalia border with Kenya, Southern parts of South Sudan, and contract in southwestern parts of Uganda (Figure 4). There was a sizeable expansion in habitat suitability for RVF in southern and central Uganda in the period 2050 and 2070 under RCP 4.5 and RCP 8.5. A large area in northern Zambia in the period 2050 was projected to expand under both RCP 4.5 and RCP 8.5 scenarios. In contrast, RVF’s habitat suitability in Rwanda was observed to contract in all future scenarios under both periods.

**Figure 3.**
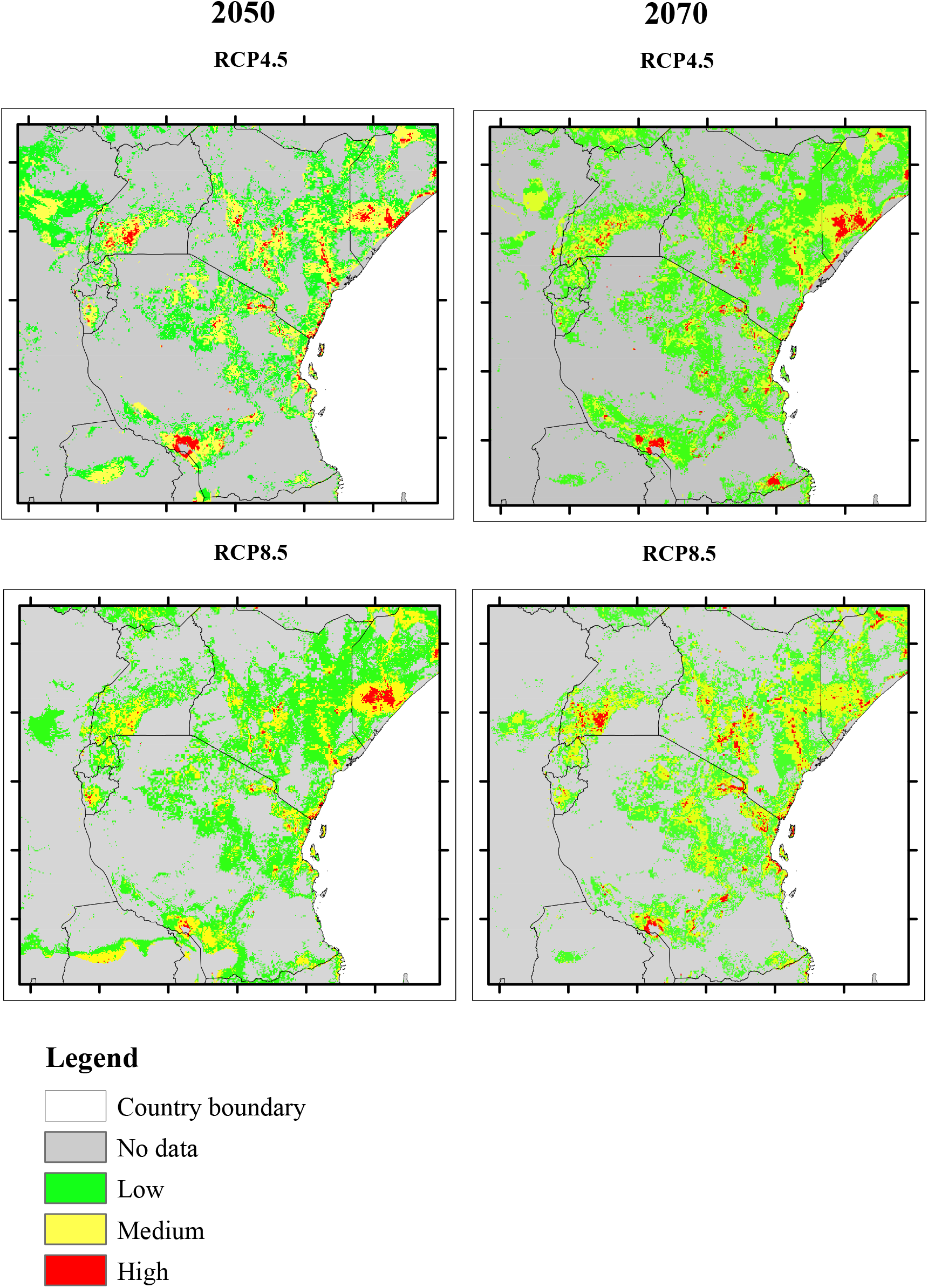
Potential climatic suitability for Rift Valley fever through time and under different RCPs.

**Figure 4.**
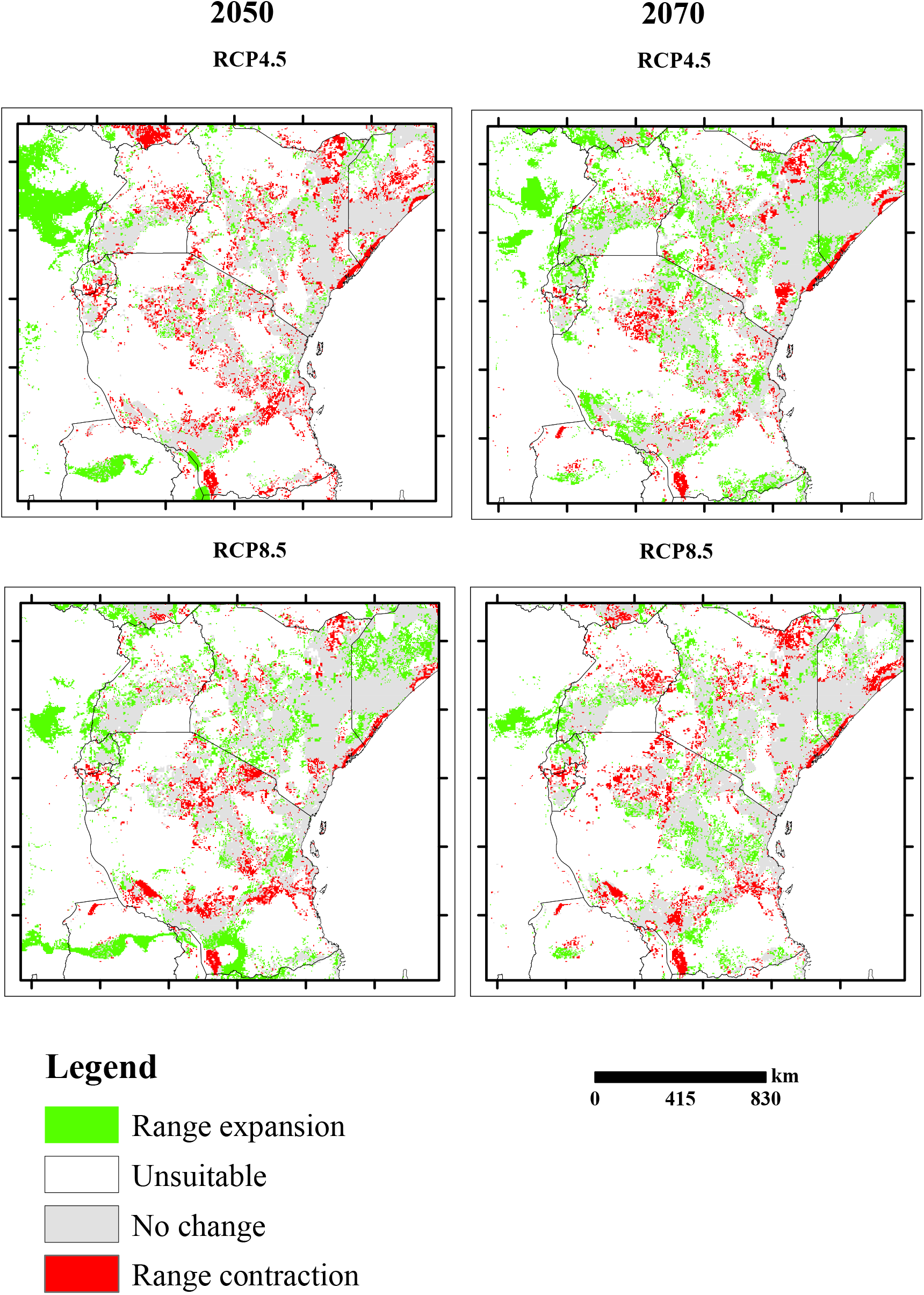
Distributional changes Rift Valley fever in East Africa, obtained by comparing binary changes between the current and future potential distribution.

## Discussion

Knowledge of zoonotic diseases’ prevalence is usually the main driving force behind evaluating how they thrive. Developing species distribution models of such diseases may help researchers quantify the magnitude, extent, and risks posed to humans, plants and animals, and their potential impacts on communities (30–31). In this study, we present the main results of MaxEnt modeling using environmental covariates and RVF outbreak cases on the disease distribution in East Africa. This is, to our knowledge, one of the most comprehensive studies about the topic in the region.

### Uncertainty of the results

Predictive habitat suitability in an ENM approach for any zoonotic is efficient as this helps to inform applied management (29). Regardless, several demerits still exist to niche modeling, for instance, low transferability in the future and the high differences between simulation results and the suitable niches for the diseases, which might lead to biased conclusions drawn from the models. Therefore, improving the simulated model transferability may generate relevant reference data for management and disease tracking and accurately predict potential niches.

Previous studies have utilized MaxEnt models that have proved to outperform other models (32). The present study models were validated by AUC statistics of the ROC, which showed high reliability. Nevertheless, few uncertainties still existed that could not be avoided in the simulations of the models, just like any other model. For instance, for some points that lacked absolute coordinates, online gazetteers and Google earth were used to geo-reference these localities using the available descriptions.

### Potential distribution and importance of variables

A set of bioclimatic variables, elevation, and animal and human population densities contribute to RVF distribution in East Africa. The variables Bio4 (temperature seasonality), land use, and population density contributed strongly to defining the distribution RVF. The variations in temperature (temperature seasonality) appeared to favor the expansion of the distributional ranges of RVF from lowlands to highlands or mountainous regions. The results are consistent with previous reports that RVF persists in the middle to low land areas in southern Africa and Eastern Africa (18) and occupies areas with dynamic temperature conditions and varying habitats ranging from wet to dry (17, 21).

The highest suitable areas for the distribution of RVF were located in eastern Kenya and southern Somalia, southern Tanzania and along the Indian Ocean coastline, and in some patches on the borders between Uganda and Kenya, and central Rwanda. These areas were observed to be low to midland areas. Livestock keeping is mainly practiced in the lowlands to midlands or plains, where grasslands are denser than forests (17, 27); as such, livestock densities were specified by the models as important variables affecting RVF distribution. Therefore, these areas are ideally suitable niches for the persistence of RVF and correspond to areas where cattle keeping is practiced with altitude ranging from 0-3000asl and with varying microclimates with low to high temperature and low to high precipitation. Consequently, from the results, we deduce that livestock densities seemingly were essential components for all RVF model distributions.

Our results agree with various zoonosis characteristics that can reappear and emerge in newly established areas (31). For instance, previous reports have shown that in the Indian Ocean coasts of E.A, the Chikungunya epidemic emerges after warm and dry conditions preceded by heavy rains (33–34). These heavy rains are observed to replenish water reserves during drought periods, and in turn, vectors breed successfully and cause risks for Chikungunya circulation (34).

### Future distributions

Future projections showed that suitable niches would expand in the 2050s and the 2070s under most climate change scenarios, while in some areas, the niches would contract. Under the RCP8.5 scenario of no mitigation of greenhouse gas emission in the 2050s and the RCP4.5 of controlled greenhouse gas emission, we observe the highest range expansion (Figure 4). We also observe RVF niches’ contraction in northern Kenya, southern Tanzania in the region bordering Zambia, the coastal areas in all scenarios. Similarly, RVF ranges are observed to decrease most under RCP4.5 in the 2050s. Under all RCP scenarios except RCP8.5 in the 2050s, parts of central and western Uganda, southern Rwanda, and northern Burundi obverse significant reduction of the suitable RVF niches. These results are comparable to a previous study by Bett et al. (17). Changes in rainfall and temperatures will likely make high elevation regions more susceptible to global warming, which will have an adverse impact on the RVF niches (15). Range expansion under climate change is typical of VBDs, which are favored by an increase in temperatures and an increase in precipitation (35–36). The West Nile virus ranges were also predicted to expand in the future in a previous study (32).

Undoubtedly, most RVF ranges in East Africa will increase due to climate change, pastoralism, human movement, anthropogenic practices, and trade in the East Africa region (37, 38, 39). Due to the alteration of rainfall and temperatures, the RVF’s vector limits are largely susceptible to global warming, which will have a considerable impact on their habitat preferences and reproduction sites’ availability (24, 35). Global climate change is projected to influence vectors, hosts, and VBDs distributions since they are all affected by temperatures (24). Therefore, VBDs will likely move polewards and to higher elevations as rapid changes in temperature drive species migrations to higher elevations (39). In addition, the gradual population growth and convention of non-agricultural areas in highlands could pose risks of RVF vectors establishment in higher elevations and latitude gradients.

The recurrence of pathogens by opportunistic host switching may persist as a significant cause of human transmittable diseases (20). The present-day niche assessment for RVF ranges and their habitat preference changes due to global climate change demonstrates concern for the public health status and further damage to livestock production. Strategies for improving public health have concentrated on enhancing monitoring in places with suspected high prevalent areas (40). With each successive outbreak, RVF has shown to be expanding; this could be attributed to the establishment of favorable climate in new areas through climate change and the introduction of vectors into new territories with favorable environments.

While climate is known to structure the ecology and distribution of species (35), other factors also come into the picture, for instance, vector abundance and distribution, land use; a sensible assumption is that climate is a significant driving force. It is estimated that RVF overburden may cause significant losses in cattle production in East Africa; however, monetary losses arising from it have not been accurately quantified and have continually risen over the years (41). To mitigate the losses brought about by VBDs, and effectively control their re-emergence, the epidemiology of the vectors and pathogens, hosts, and transmission modes for these diseases and realized niches must be understood (42, 43).

### Implications of the study

The current study that included RVF modeling in East Africa is essential in disease management, especially in regions likely to experience reoccurrence and re-emergence. Therefore, this study supports the attentiveness and management schemes of the RVF virus spread potential under climate change. Specifically, our results highlight the significance of using the disease outbreak cases to simulate the potential niches for the RVF disease in East Africa in an ENM approach, thereby revealing regions that would be otherwise overlooked when tracking the disease. The current study also highlights some areas at higher risks and prevalence of the disease, especially in Kenya and some adjacent regions in Somalia. While we cannot ascertain this from the current models, we infer that human population movement, cattle movement, and Bio4 (temperature seasonality) will play a significant role in spreading RVF in the near future (35).

Anthropogenic activities and cattle practicing, especially in densely populated areas, are likely to enhance RVF spread (37). Similarly, the vector fitness is affected directly or indirectly by temperature and increases typically with an increase in temperature, although results may differ among RVF vectors (24, 35). In addition, temperature also significantly affects vectors’ life history traits at their developmental stages, including behaviors like mating and blood-feeding (24 Approximately 30% of the total population across the East Africa region inhabits areas with suitable climatic conditions and competent mosquito vectors for the persistence and transmission of RVF (45).

The present study offers a comprehensive investigation of the effects of environmental factors linked with RVF distribution in East Africa. Our results infer that the present distribution could be due to the contemporary climate jointly with human population density and land use. Nevertheless, the absolute contributions of climate change are not apparent. Our models also indicated that climate (Bio4) had the strongest effect on the niche models developed for RVF, followed by land use and population density in that order. These results are in line with previous reports that RVF is mainly affected by environmental factors (21, 25, 29). It is not clear whether these categories’ predictions result from the locations selected for this study; however, the models were developed impartially and represent the known biology of RVF.

The forecasted increases in RVF niches will affect livelihood in E.A, particularly pastoralists who might suffer substantial losses in their livestock. Therefore, more detailed analyses are especially required to assess the localized influence of anthropogenic activities and movement within the region. We also note that the predicted contraction in RVF ranges in some areas is not guaranteed. This is primarily due to the continued urbanization and human growth, as well as the convention of the forested regions into farmlands, which might increase the potential sites for the persistence of RVF vectors. Therefore, improved surveillance in these regions is imperative to mitigate the risks that would otherwise be overlooked (22).

### Conclusions

Forecasting how future climate change will impact the transmission of VBDs has proven to be challenging. This is because the relationships between nature, human activities, and climate are complex. The current research discourses the biological response of RVF considering different climate scenarios in the current and future timeframes and its potential habitat suitability. The present climate scenario shows that RVF’s distribution is mainly in drier areas, plains, and highlights where most animal farming and pastoralism are practiced. The temperature seasonality (Bio4), land use, and population density play an important role in controlling the biogeographic patterns of RVF in the East Africa region. Thus, ENMs play an essential role in determining potential ranges suitable for RVF establishment and where the disease is most sensitive to climate change.

Although a sizeable number of studies have been utilized and developed in assessing the distribution of zoonotic diseases and the results yielded a vital contribution to the field, the present study has shown usefulness in presenting the diseases’ distribution using their actual range outbreaks. This helps to directly capture the relevant information on biotic and abiotic factors surrounding the spread of such diseases. Particularly, modeling the RVF distribution using vectors may sometimes be problematic, especially since a vector does not directly translate to RVF’s presence, and these intermediate interactions between the vectors and disease are at times less suitable descriptors of zoonotic diseases. Thus, using the disease outbreak data from the hosts may improve the species distribution of zoonotic diseases.

As a result of the increased anthropogenic activities, livestock movement, urbanization, and predicted climate change, there is a real risk that these factors might jointly enable the spread of RVF in East Africa. East Africa continues to be an endemic region for RVF, and the ongoing trade partnerships (38) could lead to the introduction of the virus to new non-endemic regions, e.g., in Uganda and Burundi. Therefore, it is vital to assess its effect on RVF distribution and spread to successive suitable areas to help design timely mitigation measures and decrease the negative impacts of future climate change. The models generated here suggested that in the future, as a result of climate change, there would be more outbreaks in new areas that were less infested by the disease, e.g., regions in Rwanda and southern Tanzania. Our present findings might help track the zoonosis in East Africa and help guide targeted management of an outbreak of the disease in marginalized and least thought regions.

## Materials and methods

### Study Area

East Africa is the eastern region of the African continent lying between the latitudes 5.2°N, 11.77°S, and longitudes 28.8°E, 41.2°E, and includes five countries: Kenya, Tanzania, Uganda, Rwanda, and Burundi. The East Africa region has a generally tropical climate. Highlands have lower average temperatures compared to areas with low elevation. There are two main rainfall seasons, the long rains season from October to December and the short rains season from March to May (46). East Africa precipitation is exceptionally heterogeneous, and this is attributed to the differing topography, maritime influence, lakes, tropical circulation, and elevation (46). Mean annual rainfall in a large part of the region falls between 800-1200 nm. East Africa’s vegetation can be classified into; forest, grassland, woodland, bushland, thicket and scrub, permanent swamp vegetation, wooded grasslands, and semi-desert (47). The region covers 3,678,394 square km, and the highest peak is Mt Kilimanjaro at 5000 m a.s.l. One feature that stands out in East Africa is the Rift Valley that cuts through the region from Kenya southward to Tanzania through Uganda. The most common animal practice is cattle herding, especially in Kenya, followed by Tanzania.

### Occurrence and environmental data

The availability of localities or occurrences is critical in ecological niche modeling when the simulation of climatically suitable areas of a target disease is intended. Therefore, we obtained the presence data (geographical coordinates) for RVFV outbreak cases during 2004-2020 from the Global Animal Disease Information System (EMPRES-i; http://empres-i.fao.org/), and previously published literature. For records that were only provided on the District level and descriptions of the RVF reported cases, we used the Online Gazetteer and Google Earth software to geo-reference them to nearest centroids. In total, we obtained 300 coordinates for RVFV cases. Duplicate records and fuzzy records were manually filtered in Microsoft Excel, followed by spatial rarefying of the remaining coordinates to exclude the correlated points within 10 by 10 km grid cells using SDMToolbox (48). Spatially correlated occurrence points would otherwise cause overfitting of the models (49). Lastly, the Excel file was converted to the “.csv” format usable in MaxEnt. The spatially rarefied points used for RVF modeling in the present study are provided in Supplementary Table 1 (Figure 1).

Bioclimatic variables for performing RVF distribution modeling were derived from Worldclim2 (http://www.worldclim.org) at 2.5arc-min (50). This data includes nineteen predictor variables with precipitation and temperature dependent information within a year. The current period was represented by bioclimatic variables spanning 1970-2000. Besides, for future periods, we utilized the global circulation models (GCM)—community climate system model version 4 (CCSM4; 51), obtained from WorldClim, to simulate the habitat suitability of RVF (period 2050 average for 2041-2060, and 2070 average for 2061-2080), at a spatial resolution of 2.5arc-min. For this GCM, we selected two representative concentration pathways (RCPs) to represent intermediate and extreme greenhouse gas emissions (RCP 4.5 and RCP 8.5), respectively, therefore adequately accounting for future climate uncertainties. These RCPs are consistent with the Intergovernmental Panel’s 5th report on climate change (4)(IPCC; 2014). Modeling climate change impacts on VBDs has broadly used the CCSM4 model (17). All the raster layers were clipped to match our study area, then converted to ASCII format in ArcGIS 10.5.

Spatially correlated variables may sometimes lead to inaccurate results of model simulations (52). Therefore, a Pearson correlation of the 19 Bioclimatic variables was implemented on ENMtools to eliminate highly correlated climatic variables at a threshold of r ≥ 0.8. Eventually, eight variables were selected for the subsequent analyses following their collinearity: Bio2 Mean Diurnal Range (Mean of monthly (max temp - min temp), Bio4 (temperature seasonality), Bio7 (Temperature Annual Range (Bio5-Bio6)), Bio8 (Mean Temperature of Wettest Quarter), Bio13 (Precipitation of Wettest Month), Bio14 (precipitation of the driest month), Bio18 (Precipitation of Warmest Quarter), Bio19 (Precipitation of Coldest Quarter) (Table 1). Further, elevation was derived from Shuttle Radar Topography Mission (http://glcf.umiacs.umd.edu). The elevation layer was used to calculate “slope” on ArcGIS 10.5. In addition, we added livestock (viz. cattle, goat, sheep) and population density layers obtained from Geo-Network with a 10 × 10 km resolution (http://www.fao.org/geonetwork/srv/en/metadata.show). These datasets were resampled to match the bioclimatic variables spatial resolution of 2.5arc-min.

### Ecological Niche modeling and validation

The present-day niches for RVF were modeled for the three periods (total of five models): The current period, the year 2050 (RCP 4.5 and 8.5), and the year 2070 (RCP 4.5 and 8.5) using MaxEnt 3.4 (53). We divided our data into a training dataset, including 80 percent and 20 percent test data, to reduce uncertainty and errors from the data. The model was run with background points of 10,000 and allowed to converge at 10^5, after 5,000 iterations. The MaxEnt models were made based on the 10-fold replicates, where cross-validation was allowed (52). Over prediction bias and model validation were performed by assessing the area under curve (AUC) stats of the Receiver Operating Characteristics (ROC) in MaxEnt (54). All averaged ASCII files were converted to TIFF format using ArcGIS 10.5.

The multivariate environmental similarity surfaces (MESS) analysis was run in MaxEnt to increase the models’ reliability for future projections (55). This function (MESS) examines the multivariate extrapolation under various climate scenarios in the future, checks for potential new climatic environments, and assesses if the future predictions are out of range, with the current projections as the baseline. In addition, distributional changes between the current and future periods were evaluated to determine the impact of climate change on RVF. To achieve this, we converted all the thresholded ASCII files into binary maps in SDMToolbox (48). For converting the ASCII files, we implemented the Maximum training and sensitivity thresholds, as suggested by Liu et al. (56). Finally, the RVF niche gains or losses were calculated by assessing the difference between the current and future suitable areas.

## Acknowledgments

The authors thank Josephat K. Saina for his tremendous assistance in the model evaluation and statistics and insights on an earlier draft of the manuscript. Much appreciation is also accorded to colleagues at the Wuhan Institute of Virology, Chinese Academy of Sciences, for their invaluable support of this project. We also wish to thank Miriam Ngarega (University of Nairobi) for her critical review of the manuscript and language improvement. We extend sincere appreciation to the committed team members at Wuhan Botanical Garden. This study was funded by the Youth Innovation Promotion Association CAS.

## Supplemental information

**Figure S1** AUC result of MaxEnt modeling.

**Figure S2** Importance of environment variables to Rift Valley fever by jackknife analysis.

**Table S1** Rift Valley fever outbreak cases used in MaxEnt Modeling.

