## Supplementary figures and images for "Present and Future Ecological Niche Modeling of Rift Valley fever in East Africa in Response to Climate Change"

### Supplemental Figure 1

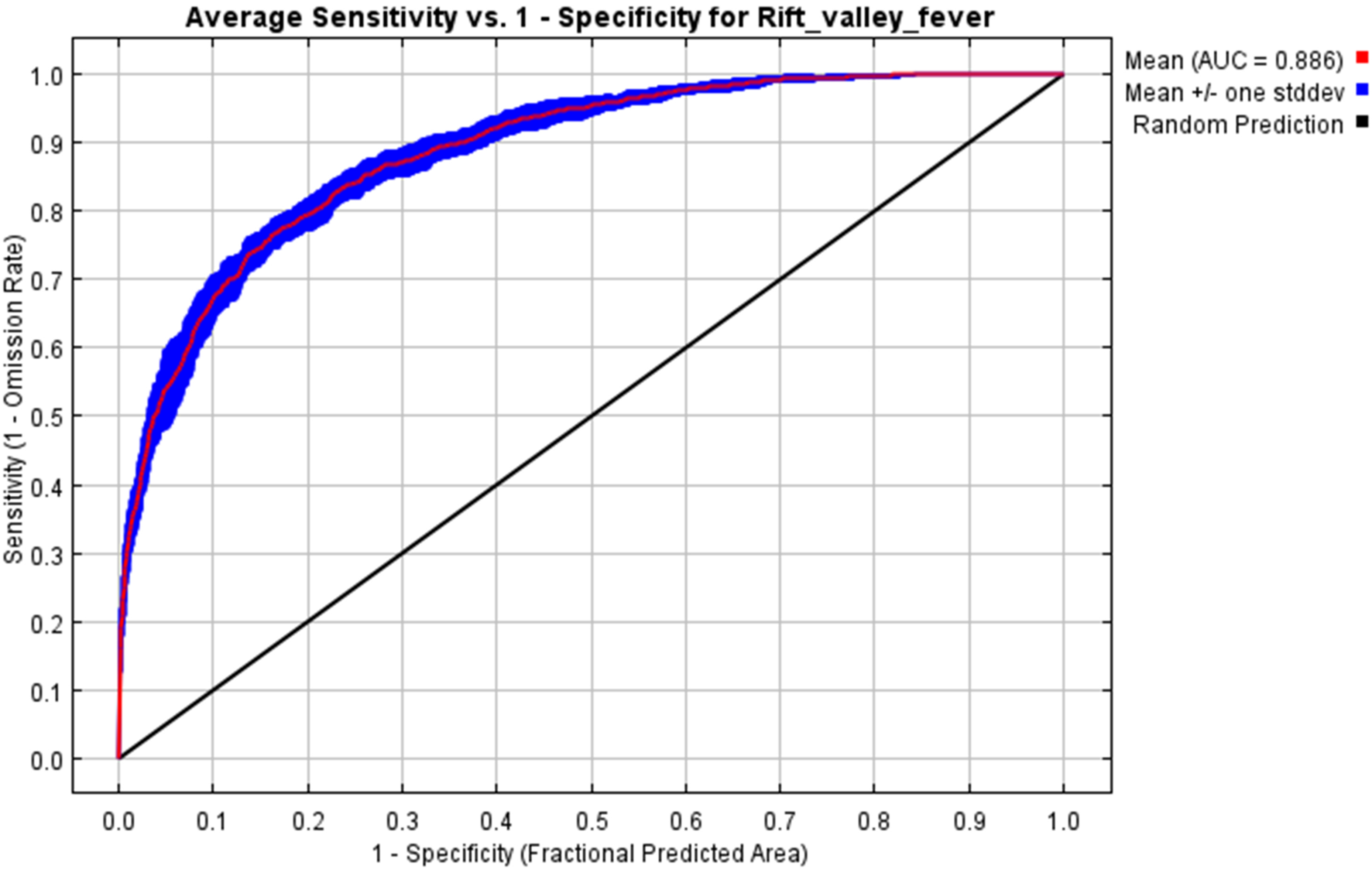

### Supplemental Figure 2

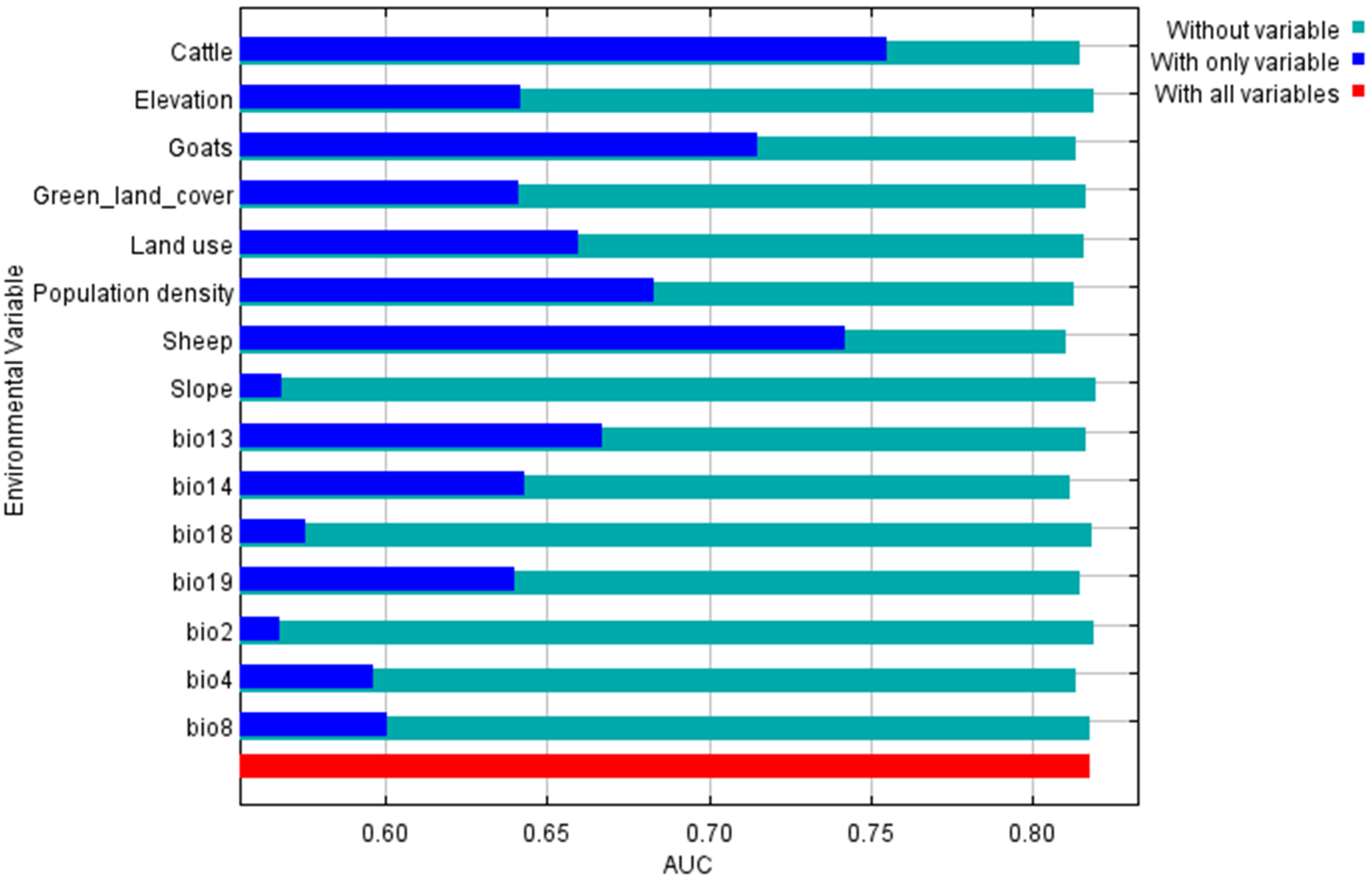
